# Complete decoupling of bacterial growth from biopolymer production through proteolytic control of enzyme levels

**DOI:** 10.1101/389809

**Authors:** Gonzalo Durante-Rodríguez, Victor de Lorenzo, Pablo I. Nikel

## Abstract

Most current methods for controlling the rate of formation of a key protein or enzyme in cell factories rely on the manipulation of target genes within the pathway. In this article, we present a novel synthetic system for post-translational regulation of protein levels, *FENIX*, which provides both independent control of the steady-state protein level and inducible accumulation of targeted proteins. The device is based on the constitutive, proteasome-dependent degradation of the target polypeptide by tagging with a short synthetic, hybrid NIa/SsrA amino acid sequence in the C-terminal domain. The protein degradation process can be reversed by activating the system *via* addition of an orthogonal inducer (e.g. 3-methylbenzoate) to the culture medium. The system was benchmarked in *Escherichia coli* by tagging two fluorescent proteins (GFP and mCherry) and further exploited for engineering poly(3-hydroxybutyrate) (PHB) accumulation completely uncoupled from bacterial growth. By tagging PhaA (3-ketoacyl-CoA thiolase, first step of the route), a dynamic metabolic switch at the acetyl-coenzyme A node was established in such a way that this metabolic precursor could be effectively directed into PHB formation upon activation of the system. The engineered *E. coli* strain reached a very high specific rate of PHB accumulation with a polymer content of ca. 72% (w/w) in glucose cultures set in the growth-decoupled mode. Thus, *FENIX* enables dynamic control of metabolic fluxes in bacterial cell factories by establishing post-translational synthetic switches in the pathway of interest.

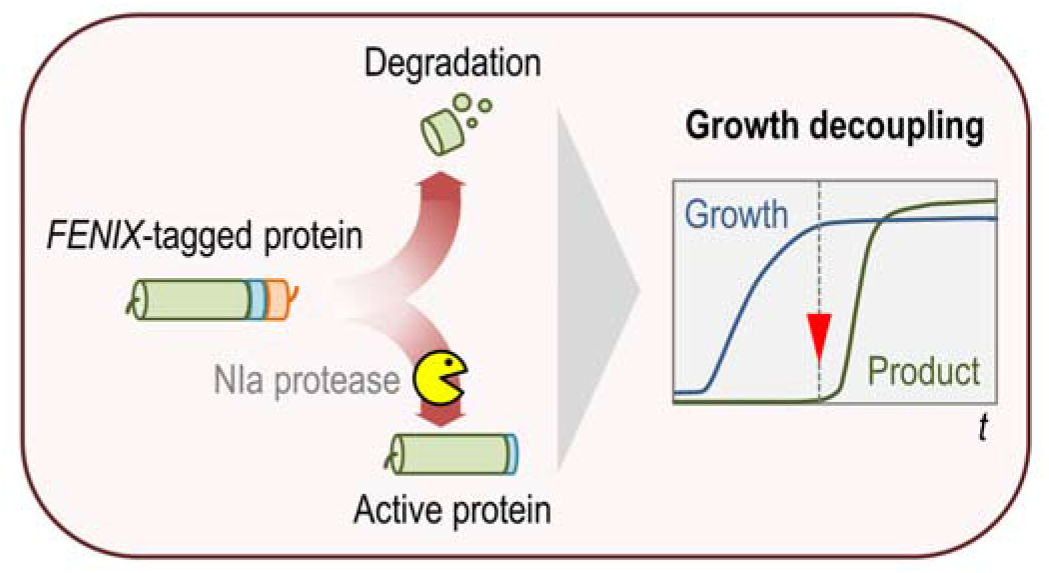
GRAPHICAL ABSTRACT.

## INTRODUCTION

One of the main challenges in contemporary metabolic engineering is to develop systems for controlling enzyme activities in a spatial-temporal fashion, leading to the highest possible catalytic output^1-2^. The problem can be tackled by manipulating genes and proteins at different levels of regulation in cell factories. Transcriptional and translational regulation mechanisms, for instance, have been studied in great detail in many biotechnologically-relevant microorganisms, and several studies describe synthetic circuits exploiting these cellular processes for practical purposes^3-6^. More recently, the adoption of CRISPR/Cas9-mediated technologies has opened up countless possibilities for targeted regulation at the gene/genome level^7-8^. The conditional and dynamic control of protein levels *in vivo*, in contrast, has received less attention thus far, and the majority of the currently available tools designed to modulate protein activity target mRNAs and protein synthesis rates (e.g. by using specific transcriptional repressors, RNA interference strategies, and riboregulators). Some synthetic devices for the tunable control of protein synthesis and degradation have been developed over the last few years^9^, e.g. systems triggered by small molecules^10-12^ or indirect degradation processes^13-15^. From a practical perspective, these strategies allow for a tight and accurate control of metabolic pathways since the transcriptional or translational regulation of the gene(s) encoding the target(s) are not altered.

Yet, as good as devices for controlling heterologous expression may be, most approaches for bioproduction of added-value compounds rely on growth-coupled biosynthesis, because constitutive expression of the genes in the target pathway is easier and simpler than inducing product accumulation after microbial growth. However, growth-coupled production severely limits product yield and productivity^16-17^. Biomass formation can consume up to 60% of the carbon source across different cultivation techniques. This situation is particularly relevant for products synthesized from precursors of central carbon metabolism that also serve as building-blocks for biomass formation. Bacterial polyhydroxyalkanoates (PHAs), biodegradable polyesters with a broad range of interesting biotechnological applications^18-19^, constitute such an example, as they are synthesized from acetyl-coenzyme A (CoA) as the main precursor^20^. PHA production in recombinant *Escherichia coli* strains has mostly exploited growth-associated polymer accumulation^21-22^, which creates a competition for acetyl-CoA between biomass formation and PHA synthesis^23^—potentially leading to metabolic imbalances that hinder high levels of product accumulation. In this context, the question at stake is whether growth and production phases could be uncoupled by repurposing natural molecular mechanisms known to control protein integrity and functionality once the cognate mRNAs have been translated.

Protein degradation in bacteria is mediated by several processes^24^. One of them is the so called *transfer-messenger RNA* (tmRNA) system, based on special RNA molecules that function both as tRNAs and mRNAs. tmRNAs form a ribonucleoprotein complex to recycle stalled ribosomes by non-stop mRNAs and tag incomplete nascent chains for degradation through the fusion of the SsrA peptide^25-26^. In *E. coli* and related bacteria, this tag sequence is recognized by the endogenous proteases ClpXP and ClpAP (that belong to the proteasome complex), which rapidly degrade the target protein. A separate proteolytic mechanism found in the prokaryotic world is the processing of viral poly-proteins. The process is mediated by enzymes that target specific amino acid sequences in otherwise very long polypeptide chains, thereby releasing functional individual proteins. One archetypal example of such poly-protein processing is based on the action of the so-called NIa protease (*nuclear inclusion protein A*)^27-28^. This enzyme was isolated from a virus of the Potyviridae family (positive-sense single-stranded RNA genome) and it has the typical structural motifs of serine proteases—although there is a cysteine residue instead of serine at the active site^29-31^. The NIa protease has been used for the proteolytic removal of both affinity tags and fusion proteins from recombinant target proteins, due to the stringent sequence specificity of the proteolytic cleavage (a mere 13 amino acid sequence)^32^.

Based on these properties, in this work we present *FENIX* (**f**unctional engineering of *SsrA/**Nla***-based flux control), a novel tool that merges the two independent degradation systems mentioned above (i.e. tmRNAs and the NIa protease), for the sake of a rapid and convenient *in vivo* control of protein activities in cell factories. To this end, a synthetic NIa/SsrA tag, which can be easily fused to the C-terminal region of any given protein *via* a single cloning step in a standardized vector, was engineered to include sequences recognized by both the protease and the proteasome. Unlike other systems for post-transcriptional regulation, the strategy relies on the constitutive degradation of the target followed by its conditional restoration. This system was instrumental to bring about an efficient decoupling of PHB accumulation from bacterial growth in recombinant *E. coli* strains by targeting a key enzyme of the PHA biosynthesis machinery.

## RESULTS AND DISCUSSION

### Rationale of *FENIX*, a synthetic post-translational control system for pathway engineering

In this work, a novel regulatory system at the post-translational level is presented that repurposes the bacterial proteasome and combines its action with the specific protease NIa, the activity of which can be externally controlled at the user’s will. While typical control devices based on proteolysis eliminate specific target proteins^33-35^, the *FENIX* system presented herein is based in just the i.e. the target is constitutively degraded by default by the endogenous proteasome until the conditional activity of the NIa protease removes the degradation signals and enables accumulation of the protein of interest (**Fig. 1**). To this end a synthetic tag sequence was designed where the recognition sequence of the potyvirus NIa protease (GESNVVVHQADER) was fused to the SsrA target sequence (AANDENYALAA) recognized by the ClpXP and ClpAP components of the bacterial proteasome^36^. The synthetic NIa/SsrA tag (GESNVVVHQADER·AANDENYALAA) can be directly fused to the C-terminal domain of virtually any protein, rendering the polypeptide sensitive to rapid degradation by the proteasome system and abolishing the protein accumulation and/or activity (**Fig. 1a**). In the presence of the NIa protease, in contrast, the proteolytic activity cleaves off the NIa/SsrA tag between the Q and A residues of the tagged polypeptide, which will then releases the SsrA target sequence from the C-terminus, thereby allowing for protein accumulation and/or enzyme activity (**Fig. 1a**).

**FIG. 1.**
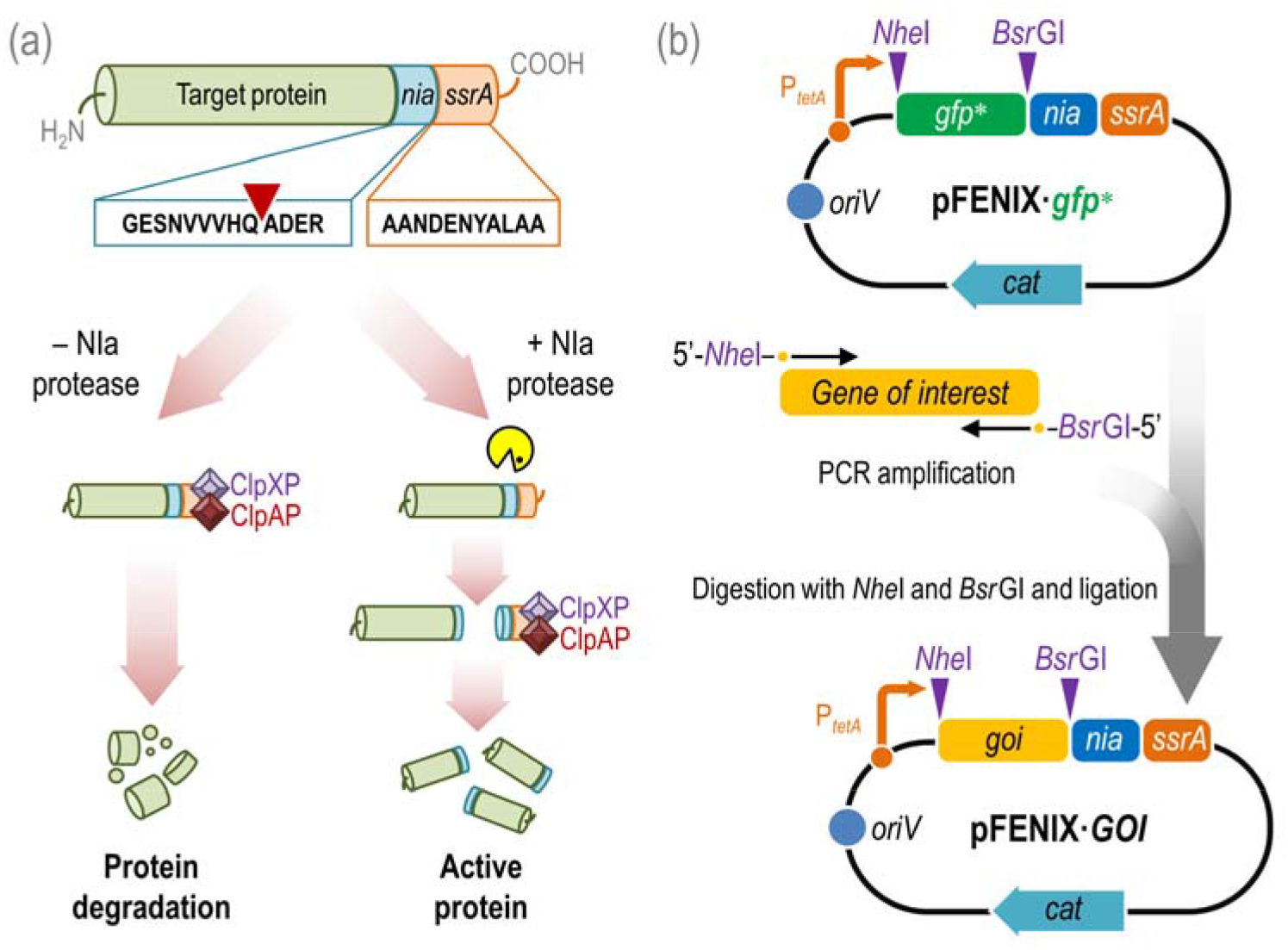
Rationale and construction of the *FENIX* system.

(a) NIa- and SsrA-dependent post-translational control of target proteins with the *FENIX* system. The gene encoding the target polypeptide is added with a synthetic *nia/ssrA* tag, resulting in a hybrid protein in which the C-terminal domain displays the GESNVVVHQADER·AANDENYALAA amino acid sequence. The SsrA tag is directly recognized by the ClpXP and ClpAP proteases of the bacterial proteasome *in vivo*, thus degrading the protein. Upon action of the specific potyvirus NIa protease (the recognition site in the synthetic *nia/ssrA* tag is indicated with an inverted red triangle in the diagram), the SsrA tag is released and the polypeptide can be accumulated. (b) FENIX plasmids for one-step cloning and tagging of individual target proteins. The gene encoding the target polypeptide (gene of interest, *goi*) is amplified by PCR with specific oligonucleotides that include *Nhe*I and *Bsr*GI restriction sites. The resulting amplicon can be directly cloned into plasmid pFENIX·*gfp*\* (which contains a *nia/ssrA* tagged version of the green fluorescent protein) upon digestion with these two restriction enzymes. In pFENIX plasmids, the expression of the *nia/ssrA-tagged* variant of the *goi* depends on the constitutive P_*tetA*_ promoter.

In order to implement this scheme, a novel set of plasmids, based on the structure set by the *Standard European Vector Architecture*^37-38^, was constructed to facilitate the direct tagging of virtually any protein sequence with the synthetic NIa/SsrA tag (**Fig. 1b**; see details in *Methods*). FENIX vectors allow for the easy exchange of the gene encoding a fluorescent protein with the coding sequence of the protein of interest upon digestion and ligation with the unique enzyme cutters *Nhe*I and *Bsr*GI (**Table 1**). The resulting FENIX plasmid will thus express a *nia/ssrA* tagged version of the gene of interest under the transcriptional control of the constitutive P_*tetA*_ promoter. An auxiliary plasmid, termed pS238·NIa, was also constructed for the regulatable expression of the gene encoding the NIa protease by placing the cognate coding sequence under the transcriptional control of the XylS/*Pm* expression system (**Table 1**), inducible upon addition of 3-methylbenzoate (3-mBz). With these plasmids at hand, we set out to calibrate the *FENIX* system as indicated in the next section.

**Table 1.**
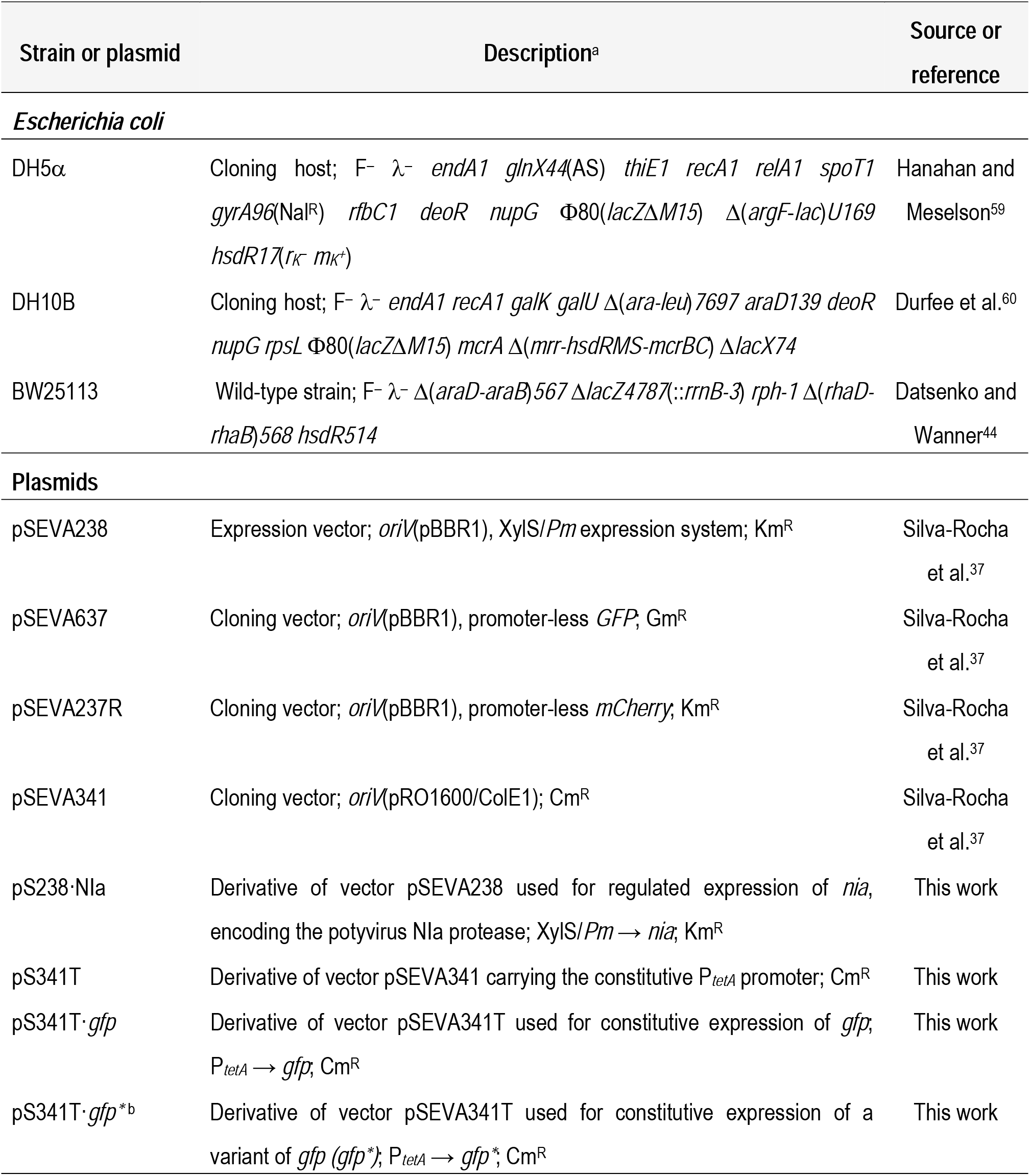
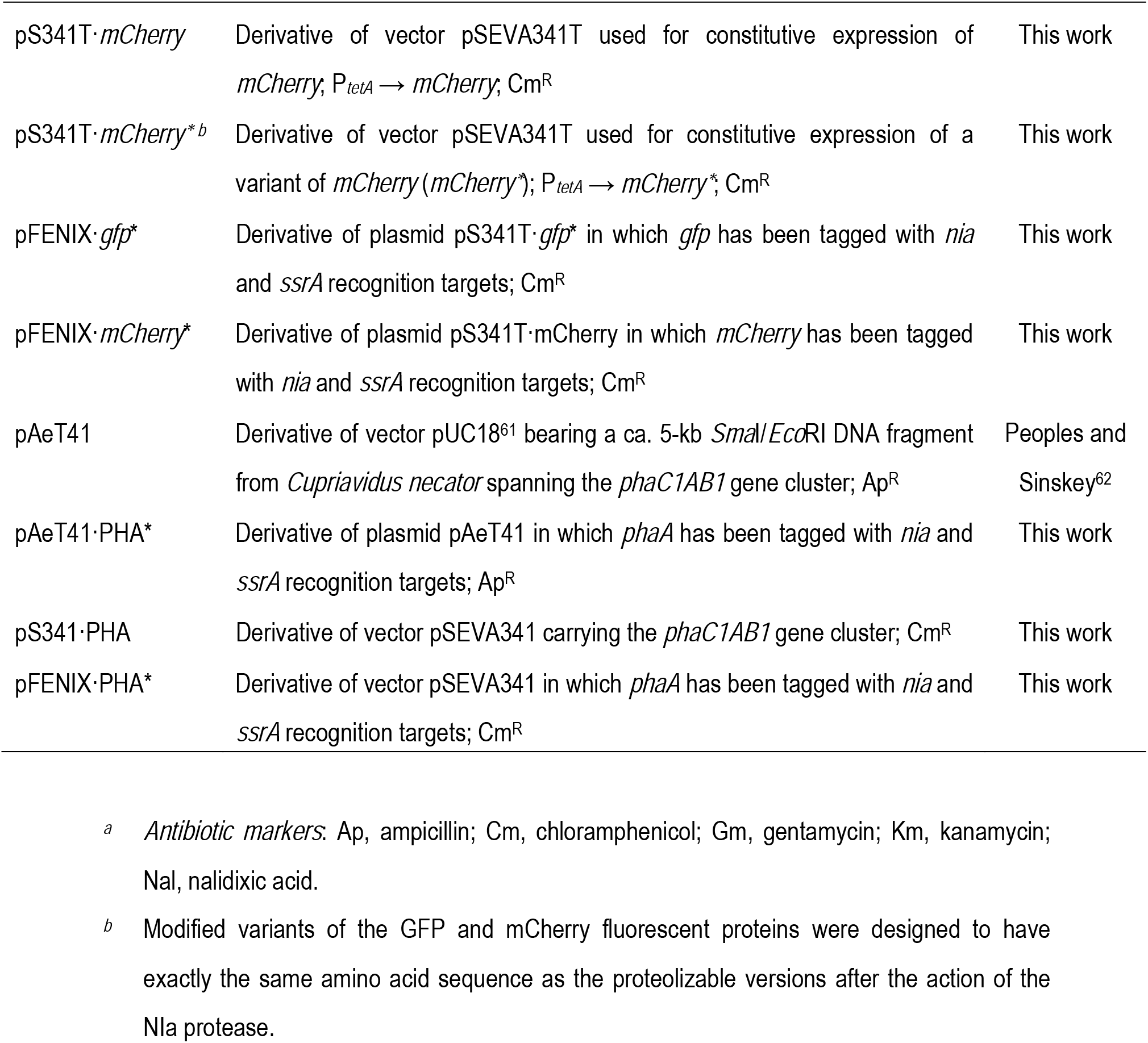
Bacterial strains and plasmids used in this study.

### The *FENIX* system enables precise control of protein accumulation in recombinant *E. coli* strains

Our first attempts at calibrating the *FENIX* system involved two fluorescent reporter proteins, the commonly-used green fluorescent protein (GFP) and the red fluorescent protein mCherry, which have been individually fused to the synthetic *nia/ssrA* tag in plasmids pFENIX·*gfp*\* and pFENIX·*mCherry*\* [**Table 1**; note that the asterisk symbol (*) indicates the addition of the synthetic NIa/SsrA tag to the corresponding polypeptide]. Each plasmid was separately transformed along with plasmid pS238·NIa in *E. coli* DH10B. When either GFP* or mCherry* are produced in *E. coli*, they will be rapidly degraded by the proteasome, i.e. no green or red fluorescence is to be seen under these conditions. Inspection of the plates in which the *E. coli* recombinants were streaked under blue light indicated that this was the case, as the colonies had no visually-detectable fluorescence (data not shown). In these strains, inducing the expression of *nia* from plasmid pS238·NIa would ultimately result in the removal of the SsrA tag, and the proteasome would no longer be able to degrade the fluorescent proteins, which could thus be detected once they accumulate in the cells at sufficient levels. To explore the kinetic properties of the *FENIX* system, these recombinant *E. coli* strains were grown in multi-well microtiter plates in LB medium with the antibiotics and additives (3-mBz) indicated in *Methods*, and bacterial growth and fluorescence (GFP or mCherry) were recorded after 24 h of incubation at 37°C (**Fig. 2**).

**FIG. 2.**
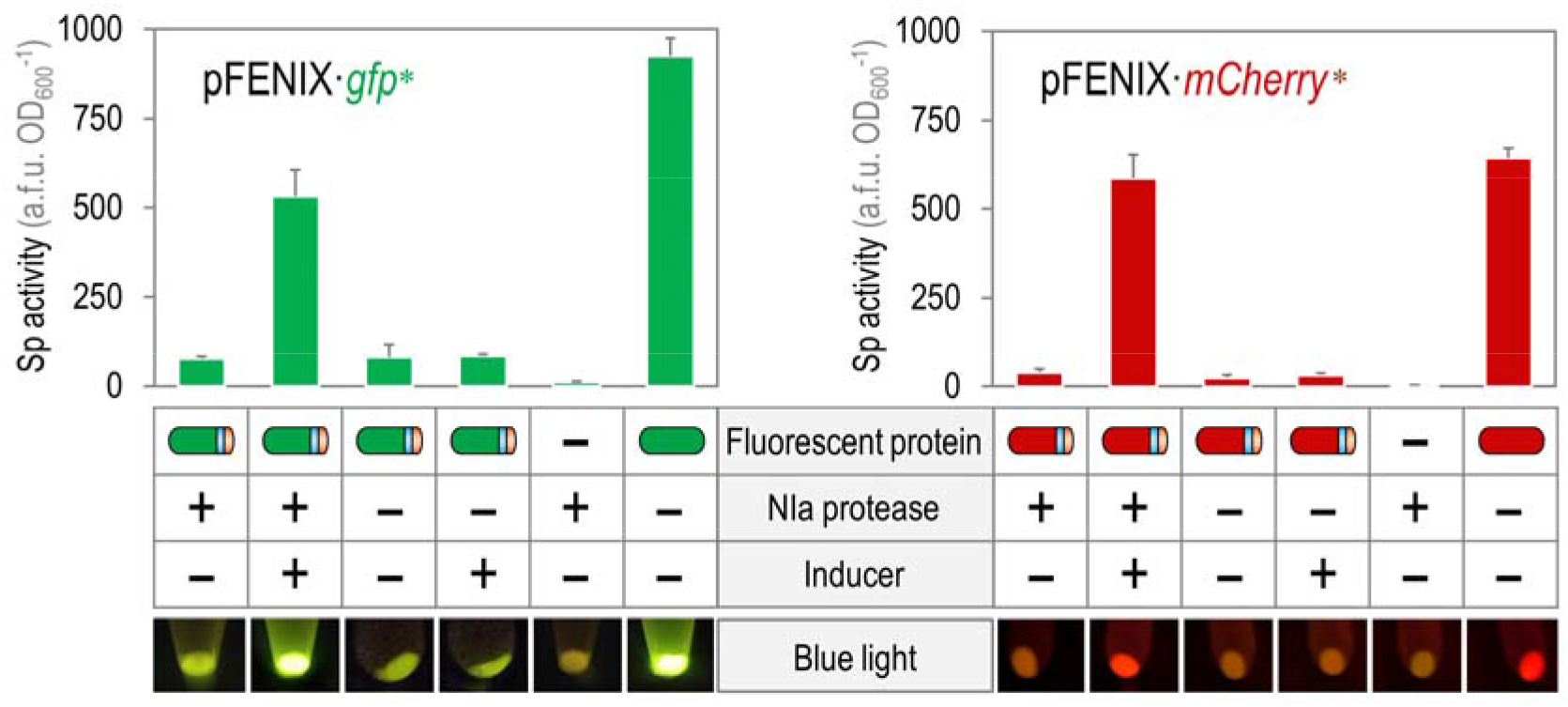
Evaluation of the FENIX system in recombinant *E. coli* using fluorescent proteins.

Plasmids pFENIX·*gfp*\* and pFENIX·*mCherry*\*, which contain the corresponding *nia/ssrA*-tagged versions of the fluorescent proteins (schematically indicated with blue and orange strips, respectively), were individually transformed into *E. coli* DH10B carrying plasmid pS238·NIa. Multiwell microtiter plates containing LB medium with the necessary antibiotics and additives (using 3-methylbenzoate at 1 mM as the inducer for *nia* expression, see *Methods* section), were inoculated with a culture of the corresponding strain previously grown overnight in LB medium with the necessary antibiotics. Cells were grown at 37°C with rotary agitation, and fluorescence and bacterial growth (expressed as the optical density measured at 600 nm, OD_600_) were recorded after 24 h. The specific (Sp) activity of the fluorescent proteins under study was calculated as the arbitrary fluorescence units (a.f.u.) normalized to the OD_600_. Each bar represents the mean value of the Sp activity ± standard deviation calculated from at least three independent experiments. The lower panel shows bacterial pellets harvested from shaken-flask cultures after 24 h of incubation under the same growth conditions indicated for the microtiter-plate cultures as observed under blue light.

The results of population-level fluorescence indicated that the qualitative behavior of the *FENIX* system was reproducible irrespective of nature of the tagged fluorescent protein. When the tagged GFP* or mCherry* proteins were exposed to the action of the NIa protease, the levels of fluorescence attained after 24 h of cultivation were comparable to those observed in the positive controls, in which the genes encoding the native (i.e. non-tagged) GFP or mCherry proteins were constitutively expressed from the P_*tetA*_ promoter (**Fig. 2**). In the case of GFP*, the final fluorescence levels were ca. 70% of those observed for GFP; for mCherry*, the fluorescence output was ca. 90% of that observed for the non-tagged version of the protein. The *FENIX* system also exhibited remarkably low levels of either GFP* or mCherry* fluorescence in the absence of 3-mBz, which indicates that the (potential) leaky expression of *nia* does not significantly affect the output fluorescence (i.e. < 10% of the fluorescence levels observed upon induction of the system in both cases)—thereby enabling tight control of protein accumulation. Moreover, and in order to explore the possible effects of the inducer of *nia* expression (3-mBz) on the behavior of the system, we also measured the specific fluorescence in cultures of *E. coli* harboring only plasmids pFENIX·*gfp*\* or pFENIX·*mCherry*\* in the presence or the absence of 3-mBz. As indicated in **Fig. 2**, the levels of specific fluorescence in either case were as low as the negative control (i.e. no fluorescent protein), irrespective of the presence of 3-mBz. These quantitative results were mirrored by the fluorescence observed in bacterial pellets of the recombinants harvested from shaken-flask cultures grown under the same conditions (**Fig. 2**, lower panel). Taken together, these results demonstrate that the *FENIX* system is functional in *E. coli* under the conditions tested, and that the proposed strategy can be established as a model for synthetic post-translational regulation. The next relevant question was to address the kinetic behavior of the system by means of flow cytometry.

### The *FENIX* system enables a precise and concerted temporal switch of protein accumulation

Since the experiments described in the preceding section analyzed the behavior of the *FENIX* system at the whole-population level, we decided to use *E. coli* DH10B transformed both with plasmid pS238·NIa and plasmid pFENIX·*gfp*\* in a set of experiments aimed at an indepth characterization of the *FENIX* system at the single-cell level (**Fig. 3**). In this case, the recombinants were grown in LB medium in shaken-flask cultures under the same culture conditions used in the experiments carried out in microtiter-plate cultures, and samples were periodically taken to analyze the levels of GFP* fluorescence by flow cytometry. At the first data point, taken at 3 h post-induction of the system by the addition of 3-mBz at 1 mM, the induced (i.e. GFP*-positive) bacterial culture behaved as a single population (i.e. characterized by a single peak in the histogram plot of cell count *versus* GFP* fluorescence; **Fig. 3a**, first panel), clearly distinguishable from the non-induced bacterial population (i.e. cultures grown in the absence of 3-mBz). This observation indicates that the operation of the *FENIX* system does not result in a mixture of sub-populations of induced and non-induced cells. The level of GFP* fluorescence rapidly increased after 5 and 8 h post-induction (**Fig. 3a**, second and third panel) and plateaued at 24 h (**Fig. 3a**, fourth panel) at fluorescence values slightly below those observed in the positive control (i.e. *E. coli* DH10B transformed with plasmid pS341T·*gfp*\*, which constitutively expresses a GFP variant displaying exactly the same amino acid sequence of the NIa/SsrA-tagged GFP *after* digestion by the NIa protease; see *Methods* for details on the construction). Interestingly, the non-induced cultures exhibited levels of GFP* fluorescence within the range of the strain used as a negative control (i.e. *E. coli* DH10B transformed with plasmid pS238·NIa) throughout the whole cultivation period—thus indicating a very low level of leakiness of the *FENIX* system in the absence of any inducer.

**FIG. 3.**
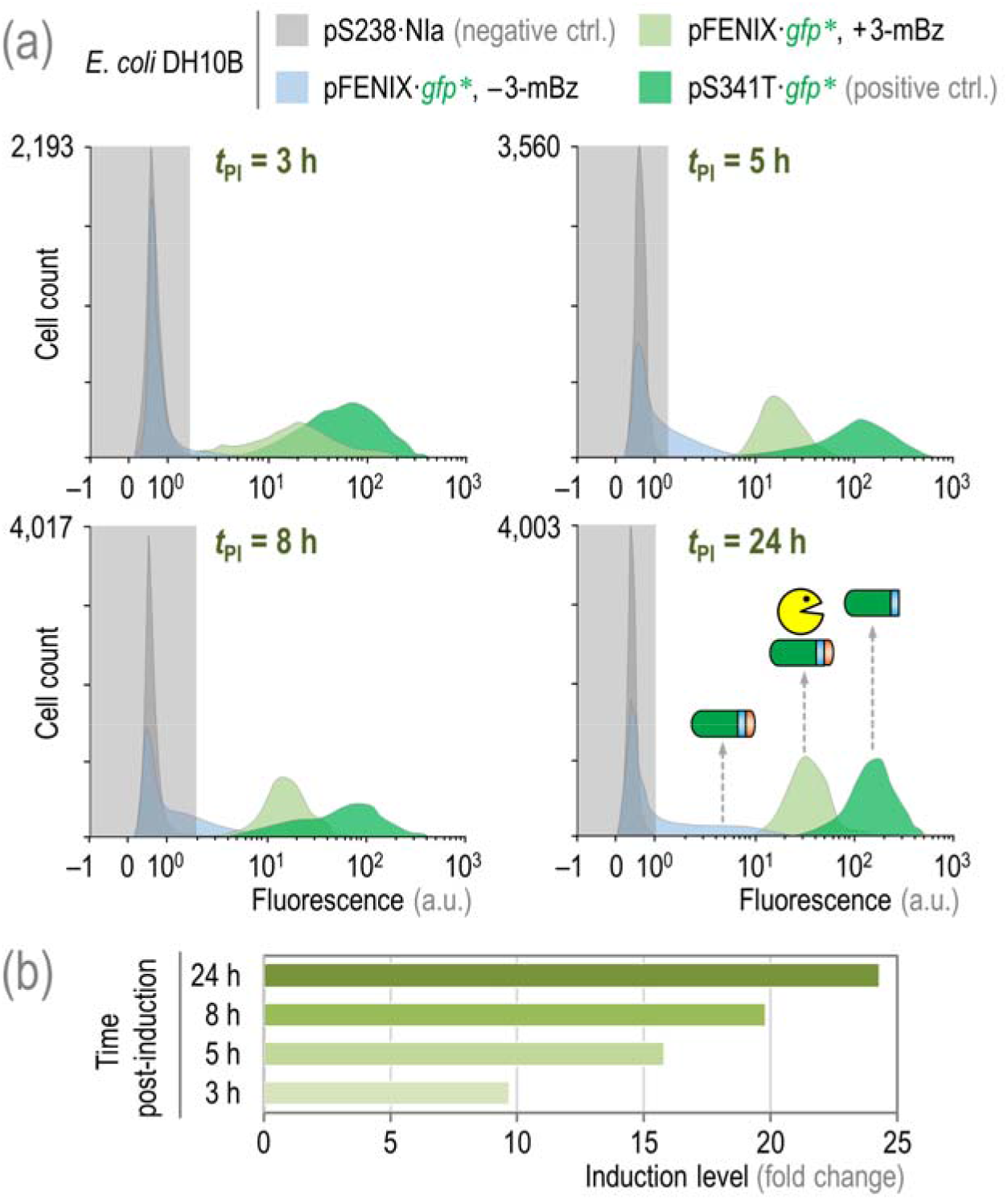
Flow cytometry analysis of the *FENIX* system.

(a) Time-lapse flow cytometry analysis of GFP* fluorescence (in arbitrary units, a.u.) in shaken-flask cultures of *E. coli* DH10B carrying the plasmids indicated. Cells were grown in LB medium at 37°C with rotary agitation with the appropriate antibiotics and additives explained in the *Methods* section, and samples were taken at selected times post-induction (*t*_pI_). The induction of the *FENIX* system was achieved by addition of 3-methylbenzoate (3-mBz) to the cultures at 1 mM at the onset of the cultivation. The light grey rectangle in each histogram plot identifies the region considered negative for the fluorescence signal (as assessed with cells carrying plasmid pS238·NIa). The structure of the *nia/ssrA-tagged* GFP and variants thereof is schematically shown in the last panel (the blue and orange strips represent the NIa and SsrA tags, respectively) along with the NIa protease (in yellow). Note that a modified version of GFP, displaying exactly the same amino acid sequence of GFP* *after* proteolysis, has been used as a positive control (ctrl.). (b) Induction levels of the *FENIX* system as calculated from flow cytometry experiments.

When the induction levels were calculated in this experiment (i.e. GFP* fluorescence in cells from induced cultures as compared to those in the non-induced control experiments), a linear increase in the fluorescence fold-change was observed over time (**Fig. 3b**). By the end of the experiment (i.e. 24 h post-induction with 3-mBz), the GFP* fluorescence levels in cultures of *E. coli* DH10B transformed both with plasmids pS238 NIa and pFENIX·*gfp*\* was 24-fold higher than those observed in the non-induced cultures of the same strain (and ca. 60-fold higher than those in cultures of *E. coli* DH10B transformed only with plasmid pS238·NIa, used as the negative control in these experiments). These results accredit the versatility of the *FENIX* system to externally control the accumulation of a target protein in a tightly regulated, and temporally coordinated fashion. Once the calibration of the system was complete, we exploited *FENIX* for tackling a longstanding problem in metabolic engineering of biopolymers as disclosed below.

### Establishing a *FENIX*-based metabolic switch for biopolymer accumulation in recombinant *E. coli* strains

*E. coli* is a suitable host for engineering biopolymer biosynthesis as it lacks the machinery needed for PHA accumulation and degradation^39^, offering the flexibility to manipulate both native and heterologous pathways for biopolymer production^40^. PHAs are ubiquitous polymers that attract increasing industrial interest as renewable, biodegradable, biocompatible, and versatile thermoplastics^41^. Poly(3-hydroxybutyrate) (PHB) is the structurally simplest and most widespread example of PHA in which the polymer is composed by C4 (i.e. 3-hydroxybutyrate) units. The archetypal PHB biosynthesis pathway of the Gram-negative bacterium *Cupriavidus necator* comprises three enzymes^42^ that use acetyl-CoA as the precursor and NADPH as the redox cofactor (**Fig. 4a**). PhaA, a 3-ketoacyl-CoA thiolase, condenses two acetyl-CoA moieties to yield 3-acetoacetyl-CoA. This intermediate is the substrate for PhaB1, a NADPH-dependent 3-acetoacetyl-CoA reductase. In the final step, (*R*)-(–)-3-hydroxybutyryl-CoA is polymerized to PHB by the PhaC1 short-chain-length PHA synthase. Expression of the *phaC1AB1* gene cluster from *C. necator* in *E. coli* results in the glucose-dependent accumulation of PHB, and several examples of metabolic engineering of biopolymer accumulation have been published over the last few decades^18-20^. Yet, the spatiotemporal control of biopolymer accumulation continues to prove challenging. On one hand, draining of acetyl-CoA away from central carbon metabolism interferes with bacterial growth if the PHB biosynthetic pathway is expressed during the active growth phase. On the other hand, acetyl-CoA is a hub metabolite in the cell, used as a precursor by a large number of metabolic pathways, and achieving precursor levels leading to high levels of PHB accumulation is inherently difficult considering the number of competing routes that also use acetyl-CoA. We hypothesized that the efficient uncoupling of bacterial growth and biopolymer accumulation could be an alternative for efficient PHB biosynthesis. Accordingly, the *FENIX* system was adapted to artificially control the level (and hence, the activity) of PhaA, the first committed step of the PHB biosynthesis pathway—and bottleneck of the entire route^43^—at the post-translational level in recombinant *E. coli* strains (**Fig. 4b**). In order to tackle this challenge, *phaA*, the second gene in the *phaC1AB1* gene cluster, was added with the synthetic *nia/ssrA* tag fragment in the 3’-end of the coding sequence (i.e. C-terminal domain of the protein) as indicated in *Methods*. The resulting engineered protein, PhaA*, would be constitutively degrade by the bacterial proteasome unless the activity of the NIa protease removes the SsrA tag from the polypeptide. On this background, the synthetic metabolic switch for controlled PHB accumulation based on the *FENIX* system was characterized as indicated in the next section.

**FIG. 4.**
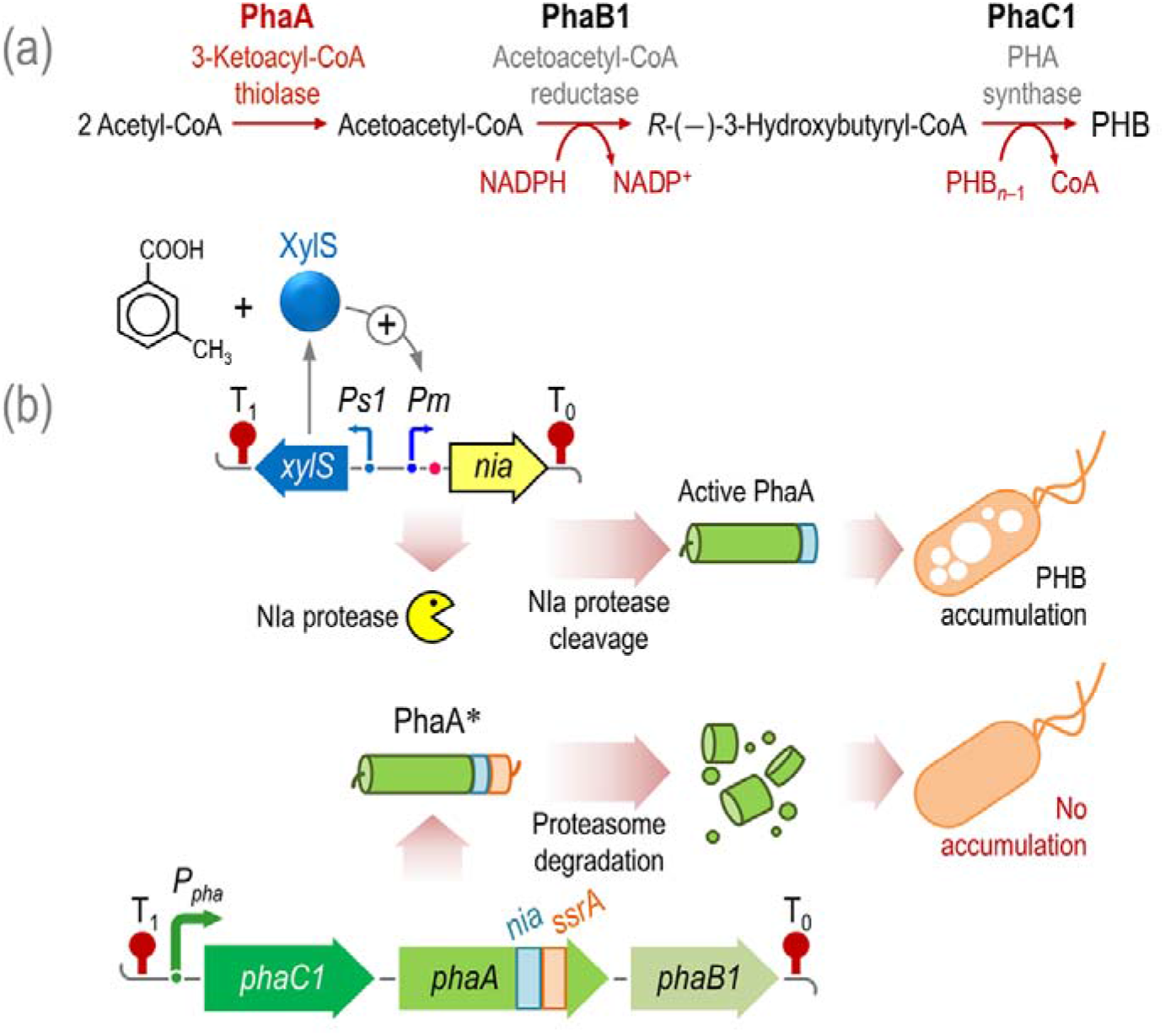
Rationale of the *FENIX*-based metabolic switch designed for controlled biopolymer.

(a) Poly(3-hydroxybutyrate) (PHB) biosynthesis pathway. Three enzymes are necessary for the *de novo* biosynthesis of PHB in *Cupriavidus necator*. 3-ketoacyl-coenzyme A (CoA) thiolase (PhaA, key step of the route as highlighted in the scheme), NADPH-dependent 3-acetoacetyl-CoA reductase (PhaB1), and PHA synthase (PhaC1). PhaA and PhaB1 catalyze the condensation of two molecules of acetyl-CoA to 3-acetoacetyl-CoA and the reduction of acetoacetyl-CoA to *R*-(-)-3-hydroxybutyryl-CoA, respectively. PhaC1 polymerizes the resulting C4 monomers into PHB, whereas one CoA-SH molecule is released per monomer. PHB is stored as water-insoluble granules in the cytoplasm of the cells. (b) Synthetic circuit based on the *FENIX* system for controlled PHB accumulation. PhaA has been earmarked with the synthetic NIa/SsrA tag in the C-terminal domain (PhaA*), thus rendering the polypeptide susceptible to proteolysis by the bacterial proteasome. Under these circumstances, no PHB is accumulated by the cells. Upon activation of the NIa protease (from a separate plasmid, in which the XylS/*Pm*-dependent expression of *nia* can be triggered by addition of 3-methylbenzoate to the culture medium), the SsrA tag is removed from the protein, the active PhaA enzyme accumulates in the cells and so does PHB. The genetic elements in this scheme are not drawn to scale. FIG. 5. Physiological and biochemical characterization of *E. coli* strains carrying the *FENIX* system tailored for controlled PHB accumulation.

### The PhaA activity can be tightly regulated by means of the *FENIX* system

*E. coli* BW25113, a well characterized wild-type strain^44^, was transformed with plasmids pS238·NIa and pFENIX·PHA* (**Table 1**). Plasmid pFENIX·PHA* expresses the *phaC1AB1* gene cluster of *C. necator* from its own constitutive promoter, and contains a variant of *phaA* fused to the *nia/ssrA-* tag sequence (**Fig. 4b**). Shaken-flask cultures of this recombinant strain were carried out in LB medium containing 30 g L^-1^ glucose, and growth parameters, PHB accumulation and the *in vitro* PhaA activity were periodically monitored over 24 h (**Fig. 5**). We first explored if the PhaA activity can be switched on by means of the *FENIX* system. In non-induced cultures (i.e. without addition of 3-mBz), the levels of 3-ketoacyl-CoA thiolase activity consistently remained below 2 μmol min^-1^ mgprotein^-1^ throughout the cultivation (**Fig. 5a**). This background thiolase activity was also detected in *E. coli* BW25113 transformed only with plasmid pS238·NIa, and can be accounted for by the endogenous ketoacyl-CoA thiolases of *E. coli* (e.g. AtoB and FadA). In contrast, when 3-mBz was added to the cultures at 1 mM, a quick and sharp increase in the *in vitro* PhaA activity was detected, reaching a 30-fold higher level at 8 h post-induction. By 24 h of cultivation, the PhaA activity in induced cultures had reached 6.1 ± 0.7 μmol min^-1^ mg_protein_^-1^. In a parallel experiment, *E. coli* BW25113/pS238·NIa was transformed either with plasmids pAeT41 or pS341·PHA, which constitutively express the native *phaC1AB1* gene cluster of *C. necator* (in the latter case, in the same vector backbone used for FENIX plasmids, i.e. pSEVA341). The *in vitro* PhaA activity was measured in 24-h cultures of these recombinant strains under the same growth conditions indicated above, in the absence of presence of 3-mBz (**Fig. 5b**). *E. coli* BW25113/pS238·NIa transformed either with plasmids pAeT41 or pS341·PHA had similarly high levels of PhaA activity irrespective of the presence of 3-mBz. In contrast, a clear difference in the thiolase activity was detected in *E. coli* BW25113 transformed both with plasmids pS238·NIa and pFENIX·PHA*. In non-induced cultures, the enzymatic activity remained at levels < 1 μmol min^-1^ mg protein^-1^ even after 24 h of cultivation, but the addition of 3-mBz triggered an 8-fold increase in PhaA activity. Moreover, the activity in the induced cultures carrying the PhaA* variant reached the highest levels among all experimental strains and conditions. The tighter control of protein accumulation afforded by the *FENIX* system thus contributes to 1.6-fold higher activity levels of the tagged enzyme as compared to the native PhaA.

**FIG. 5.**
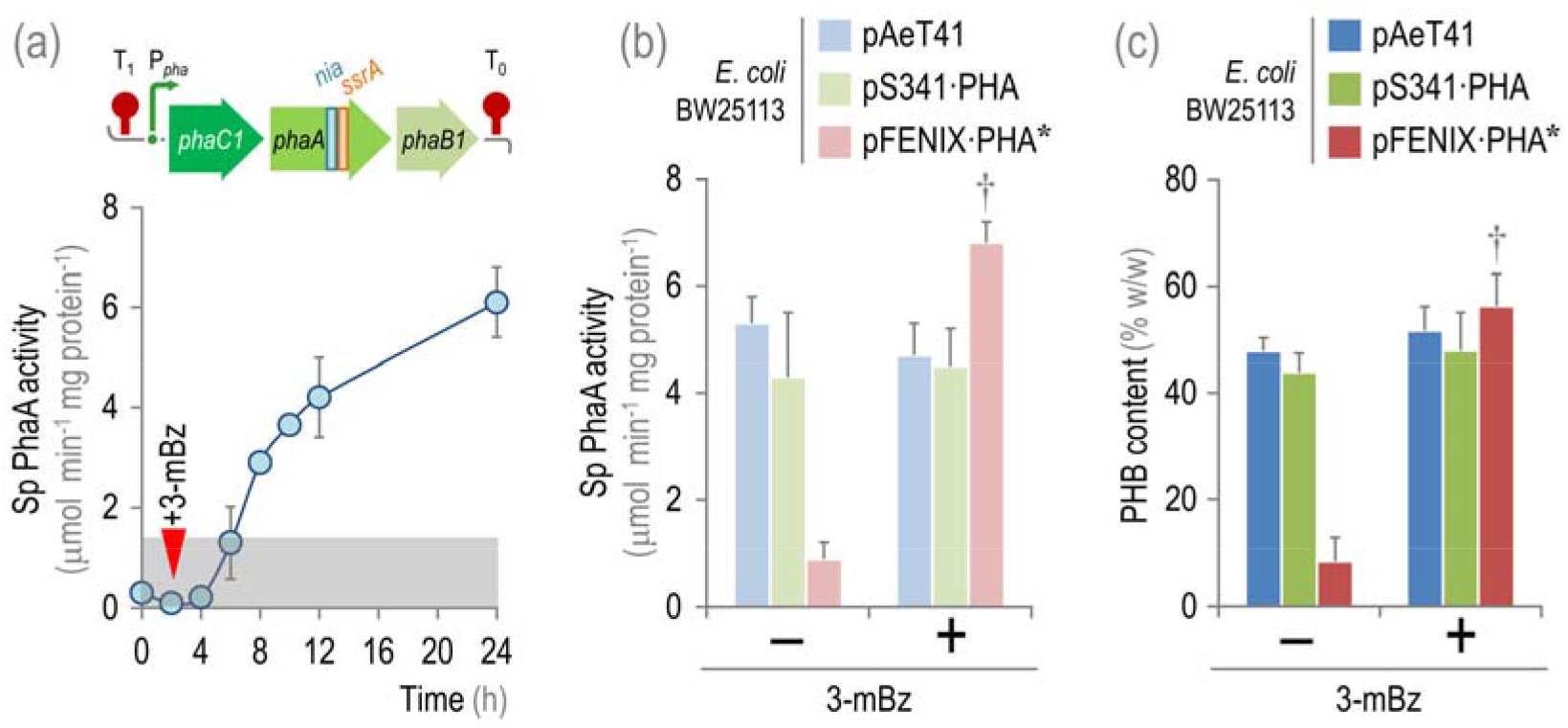
Physiological and biochemical characterization of *E. coli* strains carrying the *FENIX* system tailored for controlled PHB accumulation.

(a) *In vitro* determination of the specific (Sp) 3-ketoacyl-coenzyme A thiolase (PhaA) activity. *E. coli* BW25113 was transformed both with plasmids pS238·NIa and pFENIX·PHA* (the structure of the *nia/ssrA-tagged* variant of *phaA* in the *phaC1AB1* gene cluster of *C. necator* is schematically shown in the upper part of the figure), and the PhaA activity was periodically determined in cell-free extracts as detailed in *Methods*. The inverted red triangle indicates the addition of 3-methylbenzoate (3-mBz) at 1 mM to the culture medium; the gray bar identifies the maximum thiolase activity detected in *E. coli* BW25113 transformed only with plasmid pS238·NIa. (b) *In vitro* determination of the Sp PhaA activity and (c) PHB accumulation in *E. coli* BW25113 carrying vector pS238·NIa and the indicated plasmids. Plasmids pAeT41 and pS341 ·PHA express the native *phaC1AB1* gene cluster of *C. necator* in different vector backbones. In all plasmids used in these experiments, the expression of the *pha* gene cluster is driven by the native, constitutive P_*PHA*_ promoter. All shaken-flask cultures shown in this figure were carried out in LB medium added with 30 g L^-1^ glucose and the adequate antibiotics and additives specified in *Methods*. Each parameter is reported as the mean value ± standard deviation from duplicate measurements in at least three independent experiments. Significant differences (*P* < 0.05, as evaluated by means of the Student’s *t* test) in the pair-wise comparison of induced *versus* non-induced cultures are indicated by the † symbol.

The levels of PHB accumulation were also inspected in these cultures by means of flow cytometry and gas chromatography as indicated in *Methods*. The content of PHB in the bacterial biomass closely mirrored the levels of PhaA activity in all recombinants (**Fig. 5c**). Again, 3-mBz–induced cultures of the strain carrying the NIa/SsrA-tagged variant of PhaA exhibited the highest polymer content on a cell dry weight (CDW) basis [56.2% ± 6.1% (w/w), 7-fold higher than that in non-induced cultures] among all strains tested. Importantly, all the strains grew at similar levels (with a final biomass density of ca. 5 g_CDW_ L^-1^ at 24 h), indicating that the differences observed in PHB accumulation across the recombinants can be attributed to the dynamics of PhaA* activity brought about by the *FENIX* system and not to any effect on bacterial growth.

### The *FENIX* system enables efficient decoupling of PHB biosynthesis and bacterial growth and leads to high rates of biopolymer accumulation

In order to gain further insights into the dynamics of PHB accumulation in our recombinant *E. coli* strains in shaken-flask cultures, we carried out a thorough physiological characterization in M9 minimal medium containing 30 g L^-1^ glucose as the sole carbon source (**Fig. 6**). To this end, bacterial growth and PHB accumulation were closely monitored over 24 h in batch cultures of *E. coli* BW25113/pS238·NIa carrying either plasmid pS341 ·PHA (native PhaA) or pFENIX·PHA* (NIa/SsrA-tagged PhaA). The growth of the two strains was comparable, and the final biomass density plateaued at ca. 3.5 g_CDW_ L^-1^ (**Fig. 6a**). The trajectory of PHB accumulation, in contrast, differed between the two strains (**Fig. 6b**). In *E. coli* BW25113/pS238·NIa carrying pS341·PHA, the amount of PHB increased exponentially throughout the cultivation period (i.e. closely resembling biomass formation), whereas in the strain carrying the PhaA* variant the accumulation of PHB was clearly dissociated from bacterial growth, consistently < 5% (w/w) during the first 8 h of cultivation. Once PHB accumulation was triggered, it rapidly increased exponentially. Similarly to the observation made in LB cultures, the strain carrying the NIa/SsrA-tagged version of PhaA attained a higher PHB content in these glucose cultures [72.4% ± 1.8% (w/w), 1.3-fold higher than that in the strain expressing the native *phaC1AB1* gene cluster; **Fig. 6b**].

**FIG. 6.**
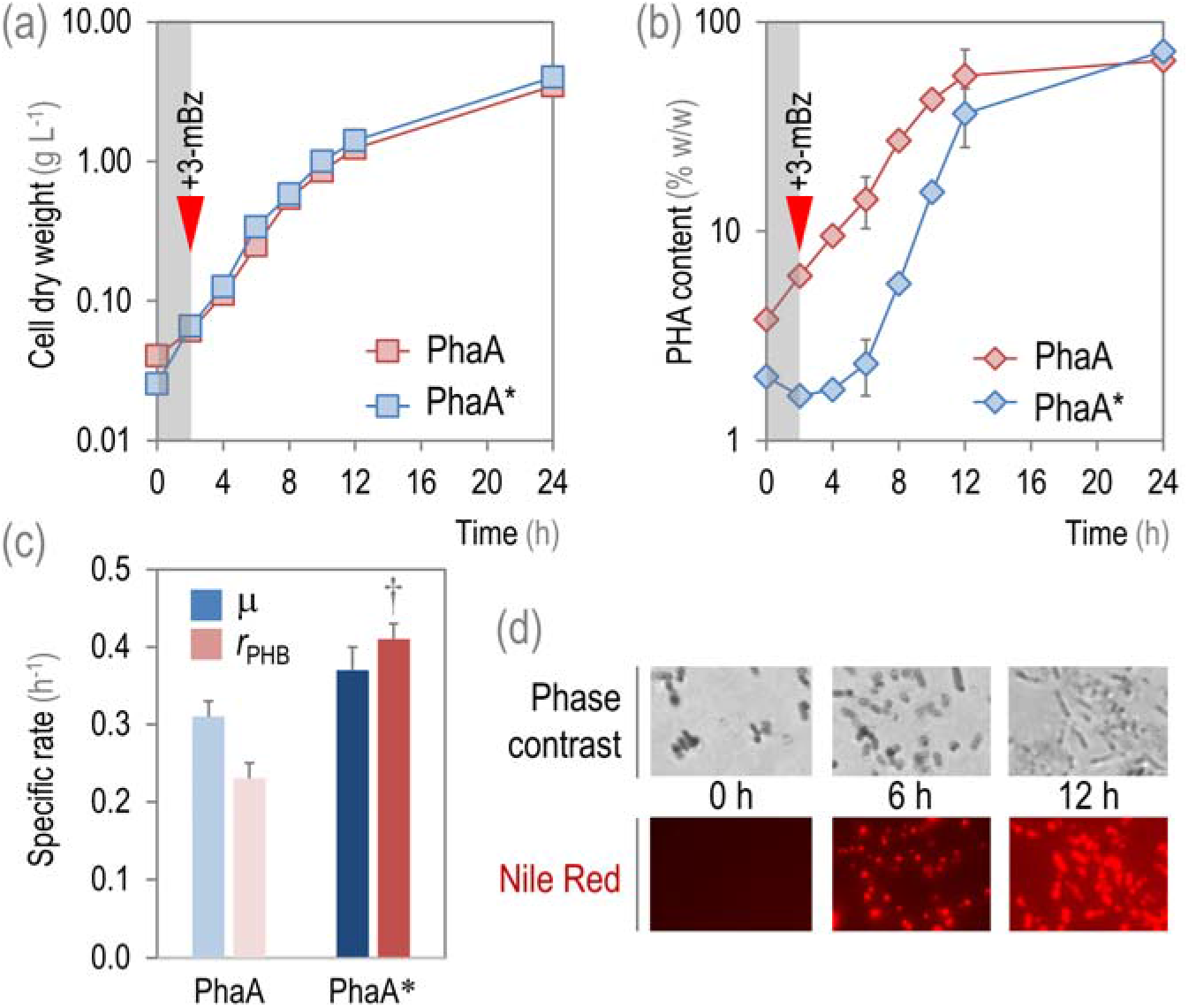
Growth and PHB accumulation by recombinant *E. coli* carrying PhaA*.

(a) Bacterial growth, expressed as the density of cell dry weight, and (b) PHB content on biomass in shaken-flask cultures of *E. coli* BW25113/pS238·NIa transformed either with plasmid pS341·PHA (expressing the native *pha* gene cluster, identified as PhaA) or pFENIX·PHA* (expressing the *nia/ssrA-tagged* variant of *phaA*, identified as PhaA*). The inverted red triangle indicates the addition of 3-methylbenzoate (3-mBz) at 1 mM to the culture medium (M9 minimal medium containing 30 g L^-1^ glucose); the gray bar also identifies the time pre-induction of the system. (c) Specific rates of bacterial growth (μ) and PHB accumulation (*r*_PHB_) in the strains under study. Significant differences (*P* < 0.05, as evaluated by means of the Student’s *t* test) in the pairwise comparison between the two strains is indicated by the † symbol. In the graphics (a-c), each parameter is reported as the mean value ± standard deviation from duplicate measurements in at least three independent experiments. (d) Qualitative assessment of PHB accumulation in samples taken from shaken-flask cultures at the indicated times and stained with the lipophilic Nile Red dye. Stained cells were observed under the microscope either under phase contrast or fluorescence as indicated in *Methods*.

Next, we assessed the specific rate of bacterial growth and biopolymer accumulation (**Fig. 6c**). The specific growth rate (μ), as inferred from the growth curves, was not significantly different between the two *E. coli* recombinants (ca. 0.3 h^-1^). However, the clear decoupling of PHB accumulation from bacterial growth in the strain carrying the PhaA* variant resulted a 2-fold higher specific rate of PHB accumulation (⊓_PHB_). Under these experimental conditions, ⊓_PHB_ = 0.41 ± 0.02 h^-1^, which is the highest reported in the literature for recombinant *E. coli* strains. The growth decoupling effect was also visually evidenced when cells were sampled from these cultures, stained with the lipophilic Nile Red dye, and observed under the fluorescence microscope (**Fig. 6d**). Upon induction of the *FENIX* system, the rapid accumulation of PHB in the recombinants could be clearly detected as the polymer granules started to fill the bacterial cytoplasm. Taken together, these results suggest that the *FENIX* system can be used as a metabolic switch operating at the acetyl-CoA metabolic node—a possibility that was explored in detail as explained below.

### Enhanced PHB accumulation mediated by PhaA* stems from flux re-wiring around the acetyl-CoA node

As indicated previously, acetyl-CoA is a metabolic hub in the cell. In the *E. coli* recombinants described in this work, a major competition occurs at this node between the PHB biosynthesis pathway and other endogenous metabolic routes. Apart from the core cell functions that use acetyl-CoA as building-block (e.g. *de novo* fatty acid synthesis), in the presence of excess glucose, *E. coli* synthesizes (and excretes) acetate from acetyl-CoA through a two-step route catalyzed by Pta (phosphotransacetylase) and AckA (acetate kinase)^45^ (**Fig. 7a**). Taking advantage of this biochemical feature, we adopted the specific rate of acetate formation and the content of acetyl-CoA as a proxy to gauge how the *FENIX* system could re-direct this metabolic precursor into a target pathway. A lower specific rate of acetate formation was detected in glucose cultures of all *E. coli* strains expressing the PHB biosynthesis pathway as compared to the control strain, transformed with the empty pSEVA341 vector (**Fig. 7b**)—consistent with a higher flux of acetyl-CoA going into PHB formation. However, *E. coli* BW25113/pS238·NIa transformed with plasmid pFENIX·PHA* had the lowest rate of acetate synthesis along all the strains tested (0.9 ± 0.1 mmol g_CDW_^-1^ h^-1^; 70% lower than that of the control strain). Interestingly, when the specific rates of glucose consumption were also determined in these cultures, no major differences were observed among all the strains (with *q*_s_ values around 7-8 mmol g_CDW_^-1^ h^-1^), indicating that the differences in acetate formation or PHB accumulation are linked to a re-routing of acetyl-CoA rather than to significant changes in the overall cell physiology.

**FIG. 7.**
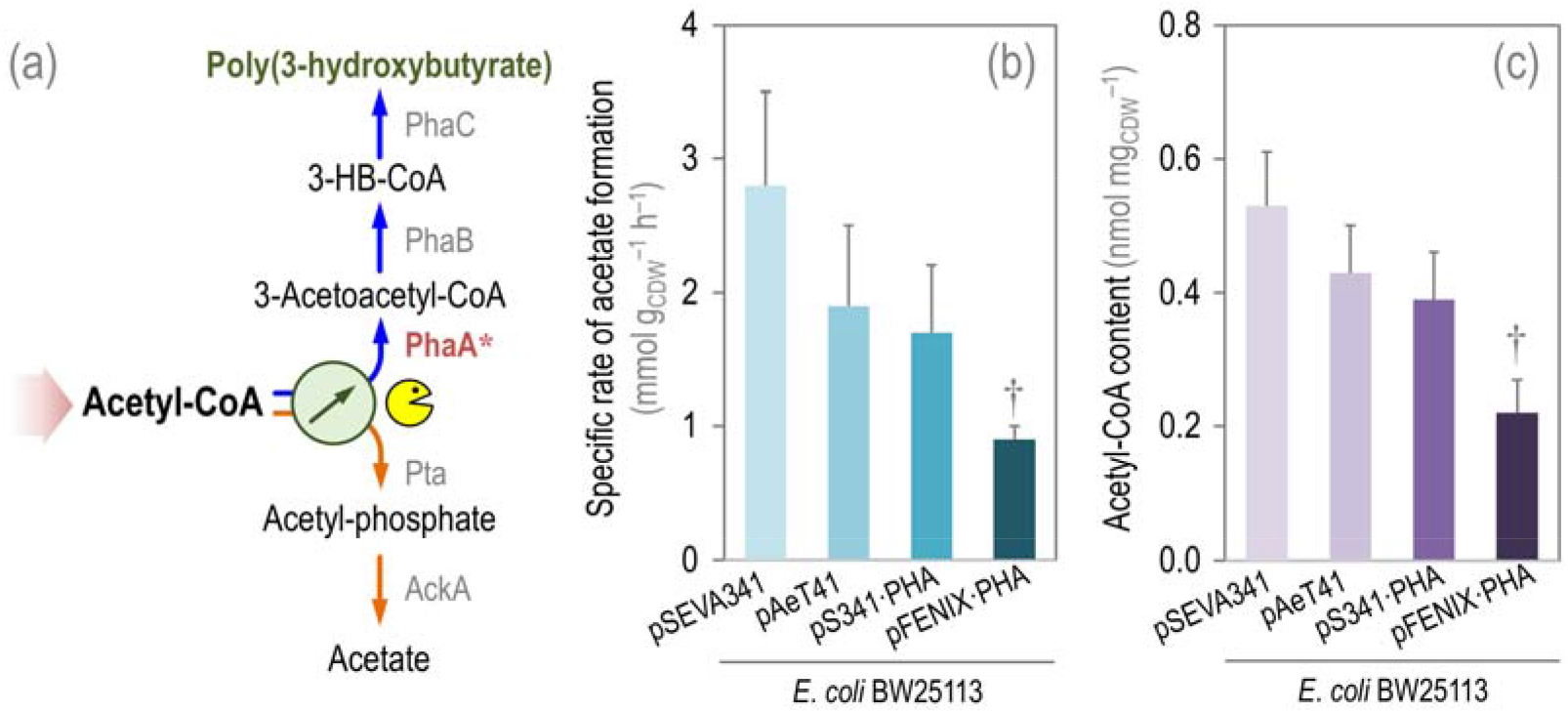
Establishing an orthogonal metabolic switch at the acetyl-CoA node based on the *FENIX* system.

(a) The acetyl-coenzyme A (CoA) metabolic node in the *E. coli* recombinants used in this study. The wide shaded arrow represents the central pathways leading to acetyl-CoA formation (i.e. glycolysis); this intermediate is used as a precursor in a myriad of metabolic reactions (not indicated in the scheme). The main sinks of acetyl-CoA are shown, namely, PHB biosynthesis or acetate formation (catalyzed by Pta, phosphotransacetylase, and AckA, acetate kinase). The NIa protease of the *FENIX* system, mediating the metabolic switch, is indicated in yellow. (b) Specific rate of acetate formation, as determined by secretion of acetate into the culture medium. (c) Intracellular content of acetyl-CoA, evaluated by LC-MS in cell extracts as explained in *Methods*. All shaken-flask cultures shown in this figure were carried out in M9 minimal medium added with 30 g L^-1^ glucose and the adequate antibiotics and additives specified in *Methods*. *E. coli* BW25113 was transformed with plasmid pS238·NIa in all cases. Each parameter is reported as the mean value ± standard deviation from duplicate measurements in at least two independent experiments. Significant differences (P < 0.05, as evaluated by means of the Student’s *t* test) in the pair-wise comparison of each recombinant against the control strain (carrying the empty pSEVA341 vector) are indicated by the † symbol. *3-HB-CoA, R*-(-)-3-hydroxybutyryl-CoA; *CDW*, cell dry weight.

The intracellular acetyl-CoA content qualitatively followed the same trend as the specific rates of acetate formation, although the values obtained for this parameter were comparable among the control strain and the *E. coli* recombinants expressing the native *phaC1AB1* gene cluster (**Fig. 7c**). Again, the tight control of the PHB biosynthesis pathway at the level of PhaA afforded by the *FENIX* system was reflected in the lowest content of acetyl-CoA among all the strains tested (0.23 ± 0.05 nmol g_CDW_^-1^)—suggesting an efficient re-routing of this metabolic precursor into PHB accumulation rather than into other metabolic sinks of acetyl-CoA. These results accredit that the *FENIX* system could be used to establish an orthogonal control over key metabolic nodes in the biochemical network, acting as a switch to re-route the fluxes around such nodes towards the formation a product of interest.

## CONCLUSION

So far, re-programming microorganisms to modify existing cell functions and to bestow bacterial cell factories with new-to-Nature tasks have largely relied on the implementation of specialized molecular biology tools—which, for the most part, tackle the issue at the genetic level of regulation. More recently, novel approaches for pathway engineering also encompass dynamic regulation of protein levels. *FENIX* exploits a hitherto unexplored feature, namely, the constitutive degradation of a target protein within a pathway, the accumulation of which can be triggered at the user’s will by addition of a cheap inducer (i.e. 3-mBz) to the culture medium. Besides the metabolic engineering application discussed in the present study (i.e. biopolymer accumulation in recombinant *E. coli* strains by targeting PhaA, the first enzymatic activity of the pathway), the *FENIX* system affords more complex pathway engineering approaches in which the formation of multiple proteins within different domains of the metabolic network can be externally controlled. The tight post-translational regulation of the system enables product titers that would be difficult to achieve by merely manipulating the level of expression of the cognate genes. Moreover, and considering the dynamic response of FENIX-tagged proteins accumulation, the system would also allow for the expression of highly toxic proteins or enzymes. These scenarios are currently under exploration in our laboratory and may lead to the development of better strategies to manipulate central and peripheral pathways to enhance the production of biochemicals and other molecules of industrial interest.

## METHODS

### Bacterial strains and cultivation conditions

The *E. coli* strains and plasmids used in this study are listed in **Table 1**. *E. coli* was grown at 37°C in LB medium^46^ or in M9 minimal medium^47^ added with glucose (30 g L^-1^) as the sole carbon source. For solidified culture media, 1.5% (w/v) agar was used. Shaken-flask cultivations were routinely carried out in an air incubator with orbital shaking at 200 rpm. Aerobic cultures were set by using a 1:10 culture medium-to-flask volume ratio. Antibiotics were added to the cultures where appropriate at the following final concentrations: ampicillin (Ap, 150 mg L^-1^), chloramphenicol (Cm, 30 mg L^-1^), and kanamycin (Km, 50 mg L^-1^).

### General molecular biology techniques

Recombinant DNA techniques were carried out by following well established methods^48^. Plasmid DNA was prepared from *E. coli* recombinants with a High-Pure plasmid isolation kit (Roche Applied Science). DNA fragments were purified from agarose gels with the Gene-Clean Turbo kit (Q-BIOgene). Oligonucleotides were purchased from Sigma-Aldrich Co. The identity of all cloned inserts and DNA fragments was confirmed by DNA sequencing through an ABI Prism 377 automated DNA sequencer (Applied Biosystems Inc.). Transformation of *E. coli* cells with plasmids was routinely carried out by means of the RbCl method or by electroporation^48^ (Gene Pulser, Bio-Rad).

### Design and construction of *FENIX* plasmids carrying proteolizable versions of GFP and mCherry

The general strategy for the assembly of FENIX plasmids is indicated in **Fig. 1b**. In all the constructs described in this article, the asterisk symbol (*) indicates that the corresponding gene has been added with a synthetic *nia/ssrA* tag. The starting point was the creation of plasmids pFENIX·*gfp*\* and pFENIX·*mCherry*\* as follows: the *nia/ssrA* tag was firstly assembled using the synthetic oligonucleotides 5’-*nia/ssrA*·*Bsr*GI (5’-GAG C<u>TG TAC A</u>AG GGT GAA AGC AAC GTG gtg gtg cat cag gcg gat gaa cgc gca gca aac gac gaa aac-3’; an engineered *Bsr*GI site, not present in SEVA vectors^37^, is underlined) and 3’-*nia/ssrA·Hin*dIII (5’-CCC <u>AAG CTT</u> TTA AGC TGC TAA AGC GTA gtt ttc gtc gtt tgc tgc gcg ttc atc cgc ctg atg cac cac-3’; an engineered *Hin*dIII site is underlined). The 42-bp long DNA sequence indicated in lowercase letters in these two oligonucleotides was used as an overlapping extension for sewing PCR, and the whole 89-bp long DNA fragment spanning the synthetic *nia/ssrA* tag was amplified with *Pfu* DNA polymerase (Promega) as per the manufacturer’s instructions. Plasmid pS341T was constructed by cloning the P_*tetA*_ promoter (a medium-strength constitutive promoter in Gram-negative bacteria in the absence of the TetR negative regulator^49^) between the PacI and EcoRI restriction targets of vector pSEVA341, and a *Nhe*I restriction target, not present in SEVA vectors, was added to the construct to facilitate further cloning. Plasmid pS341T·*mCherry* was constructed by placing the gene encoding the red fluorescent protein mCherry under control of the P**tetA** promoter as a *Xho*I/*Hin*dIII fragment obtained from vector pSEVA237R, and a *Bsr*GI restriction target was added upstream the *mCherry* coding sequence by PCR. The resulting pS341T·*mCherry* plasmid was further engineered to include the synthetic *nia/ssrA* tag by means of sewing PCR. The tag was directly cloned as a *Bsr*GI/*Hin*dIII fragment downstream the *mCherry* gene, thus giving rise to pFENIX·*mCherry*\* (**Table 1**). The same procedure was repeated with the gene encoding GFP, yielding pFENIX·*gfp*\* (**Table 1**). Both plasmids were used to calibrate the FENIX system, and they allow for the easy construction of a proteolizable version of virtually any protein by a direct cloning step of the corresponding gene of interest into the *Nhe*I and *Bsr*GI restriction sites that flank the fluorescent protein coding sequence.

Two expression vectors were also constructed as positive controls of the FENIX system. In order to stablish a direct comparison between the fluorescence originated by the engineered GFP* or mCherry* fluorescent proteins after proteolysis, we designed and created a version of these two proteins that have the same amino acid sequence as the proteolizable variants *after* digestion by the NIa protease. Plasmid pS341T·*mCherry*\*, encoding such an engineered mCherry protein, was constructed by amplifying the *mCherry* gene plus the short sequence of the *nia* target that remains after protease digestion using oligonucleotides 5’-mCherry·*Nhe*I (5’-CAC AGG AGG <u>GCT AGC</u> ATG GTG AG-3’; an engineered *Nhe*I site is underlined) and 3’-*mCherry*·*Hin*dIII (5’-GGG <u>AAG CTT</u> TTA CTG ATG CAC CAC CAC GTT GCT TTC-3’; an engineered *Hin*dIII site is underlined) by using plasmid pFENIX·*mCherry*\* as the template. The resulting amplicon, which spans the sequence encoding the mCherry protein after proteolysis, was restricted with the enzymes indicated and cloned into the *Nhe*I/*Hin*dIII-digested pS341T vector, thereby obtaining plasmid pS341T·*mCherry*\* (**Table 1**). The same procedure was repeated for GFP, yielding plasmid pS341T·*gfp*\* (**Table 1**).

### Construction of plasmid pFENIX·PHA* for post-translational control of PHB accumulation in recombinant *E. coli* strains

Since *phaA* lies in the middle of the *pha* gene cluster of *C. necator*, the strategy used for tagging this gene was slightly different as the one described above for single-gene targets. In this case, the synthetic *nia/ssrA* tag was firstly added to *phaA* by overlapping PCR. Two individual DNA fragments upstream and downstream with respect to the *STOP* codon of *phaA* were amplified by PCR using oligonucleotides (i) *5’-phaA·Bgl*II (5’-CAC GCG GCA <u>AGA TCT</u> CGC AGA CC-3’; an engineered *Bgl*II site is underlined) and *3’-phaA·nia* (5’-cgt cgt ttg ctg cgc gtt cat ccg cct gat gca cca cca cgt tgc ttt cac cTT TGC GCT CGA CTG CCA GCG C-3’) for the upstream fragment (2,462 bp) and (ii) *5’-phaA·nia* (5’-gca tca ggc gga tga acg cgc agc aaa cga cga aaa cta cgc ttt agc agc tTA AGG AAG GGG TTT TCC GGG GC-3’) and 3’-phaA·EcoRI (5’-GAC CAT GAT TAC <u>GAA TTC</u> TTC TGA ATC CAT G-3’; an engineered *Eco*RI site is underlined) for the downstream fragment (1,398 bp). Both amplicons were used to construct a DNA fragment spanning *phaA* and the synthetic *nia/ssrA* tag by sewing PCR using the overlapping sequences in the oligonucleotides *5’-phaA·nia* and 3’-*phaA·nia* (indicated in lowercase letters). This DNA fragment was cloned into the *Bg*lII/*Eco*RI-digested plasmid pAET41, obtaining plasmid pAeT41·PHA*, in which the native *phaA* sequence has been exchanged by the *nia/ssrA* tagged version of the same gene. Plasmid pAeT41·PHA* was then used as the template for a PCR amplification of the engineered *pha* gene cluster by using oligonucleotides 5’-PHA·*Bam*HI (5’-AGA <u>GGA TCC</u> GGA CTC AAA TGT CTC GGA ATC GCT G-3’; an engineered *Bam*HI site is underlined) and 3’-PHA·*Eco*RI (5’-GC<u>G AAT TC</u>C ACC GCA ATA CGC GGG CGC CAG-3’; an engineered *Eco*RI site is underlined). The resulting amplicon (4,292 bp) was digested with *Bam*HI and *Eco*RI and cloned into the same restriction sites of vector pSEVA341, resulting in plasmid pFENIX·PHA*. To test PHB accumulation using a comparable vector system, plasmid pS341·PHA was constructed as follows. The native *pha* gene cluster was amplified by PCR from plasmid pAeT41 as the template using oligonucleotides 5’-PHA·*Bam*HI and 3’-PHA·*Eco*RI. The resulting DNA fragment (4,220 bp) was digested with *Bam*HI and *Eco*RI and cloned into the same restriction sites of vector pSEVA341, resulting in plasmid pS341·PHA. *E. coli* BW25113 was transformed with plasmid pS238 NIa and either pS341·PHA or pFENIX·PHA*, and tested for PHB accumulation as indicated below.

### Flow cytometry evaluation of the *FENIX* system

Single-cell fluorescence was analyzed with a MACSQuant™ VYB cytometer (Miltenyi Biotec GmbH). GFP was excited at 488 nm, and the fluorescence signal was recovered with a 525/40 nm band pass filter. Cells were harvested at different time points as indicated in the text, and at least 15,000 events were analyzed for every aliquot. The GFP signal was quantified under the experimental conditions tested by firstly gating the cells in a side scatter against forward scatter plot, and then the GFP-associated fluorescence was recorded in the FL1 channel (515-545 nm). Data processing was performed using the FlowJo™ software as described elsewhere^50^.

### *In vitro* quantification of the PhaA activity

Cell-free extracts were obtained from bacteria harvested by centrifugation (4,000×*g* at 4°C for 10 min). Cell pellets were resuspended in 1 mL of a lysis buffer containing 10 mM Tris·HCl (pH = 8.1), 1 mM EDTA, 10 mM β-mercaptoethanol, 20% (v/v) glycerol, and 0.2 mM phenylmethylsulphonylfluoride, and lysed as described elsewhere^51^. The lysate was clarified by centrifugation (4°C, 10 min at 8,000×*g*) and the resulting supernatant was used for enzyme assays. The total protein concentration was assessed by means of the Bradford method with a kit from BioRad Laboratories, Inc. (USA), and crystalline bovine serum albumin as standard. *In vitro* quantification of the specific 3-ketoacyl-CoA thiolase activity in the thiolysis direction was conducted according to Palmer et al.^52^ and Slater et al.^42^, with the following modifications. The assay mixture (1 mL) contained 62.4 mM Tris·HCl (pH = 8.1), 50 mM MgCl2, 62.5 μM CoA, and 62.5 μM acetoacetyl-CoA. The reaction was initiated by addition of cell-free extract, and the disappearance of acetoacetyl-CoA was measured over time at 30°C (using s_304_ = 16.9×10^3^ M^-1^ cm^-1^ as the extinction coefficient for 3-acetoacetyl-CoA). The actual acetoacetyl-CoA was routinely quantified prior to the assay in a buffer containing 62.4 mM Tris·HCl (pH = 8.1) and 50 mM MgCl2. One enzyme unit is defined as the amount of enzyme catalyzing the conversion of 1 μmol of substrate per min at 30°C.

### PHB quantification

The intracellular polymer content in *E. coli* was quantitatively assessed by flow cytometry by using a slight modification of the protocol of Tyo et al.^53^ and Martínez-García et al.^54^ Cultures were promptly cooled to 4°C by placing them in an ice bath for 15 min. Cells were harvested by centrifugation (5 min, 5,000×*g*, 4°C), resuspended to an OD_600_ of 0.4 in cold TES buffer [10 mM Tris·HCl (pH = 7.5), 2.5 mM EDTA, and 10% (w/v) sucrose], and incubated on ice for 15 min. Bacteria were recovered by centrifugation as explained above, and resuspended in the same volume of cold 1 mM MgCl2. A 1-ml aliquot of this bacterial suspension was added with 3μL of a 1 mg mL^-1^ Nile Red [9-diethylamino-5H-benzo(α)phenoxazine-5-one] solution in DMSO and incubated in the dark at 4°C for 30 min. Flow cytometry was carried out in a MACSQuant™ VYB cytometer (Miltenyi Biotec GmbH). Cells were excited at 488 nm with a diode-pumped solid-state laser, and the Nile Red fluorescence at 585 nm was detected with a 610 nm long band-pass filter. The analysis was done on at least 50,000 cells and the results were analyzed with the built-in MACSQuantify™ software 2.5 (Miltenyi Biotec). The geometric mean of fluorescence in each sample was correlated to PHB content (expressed as a percentage) through a calibration curve. PHB accumulation was double-checked in selected samples by acid-catalyzed methanolysis of freeze-dried biomass and detection of the resulting methyl-esters of 3-hydroxybutyric acid by gas chromatography^23^, ^55-56^.

For microscopic visualization of PHB accumulation^57^, cells harvested from shaken-flask cultures were washed once with cold TES buffer, re-suspended in 1 mL of the same buffer to an OD_600_ of 0.4, and stained with Nile Red as indicated for the flow cytometry experiments. Aliquots of the treated cell suspension were washed once with TES buffer, immediately lay in a microscope slide, and covered with a glass cover slip (to protect the stained cells from immersion oil). Images were obtained using an Axio Imager Z2 microscope (Carl Zeiss), equipped with the scanning platform Metafer 4 and CoolCube 1 camera (MetaSystems) under a 1,000× magnification. Under these conditions, PHB granules stained with Nile Red fluoresced bright orange, with individual granules often visible within the cells.

### Other analytical techniques

Residual glucose and acetate concentrations were determined in culture supernatants using enzymatic kits (R-Biopharm AG), essentially as per the manufacturer’s instructions. Control mock assays were made by spiking M9 minimal medium with different amounts of the metabolite under examination. Metabolite yields and kinetic culture parameters were analytically calculated from the raw growth data as described elsewhere^51^. The intracellular content of acetyl-CoA was determined by liquid chromatography coupled to mass spectrometry as indicated by Pflüger-Grau et al.^58^.

### Statistical analysis

All reported experiments were independently repeated at least three times (as indicated in the figure legends), and mean values of the corresponding parameter and standard deviation is presented. The significance of differences when comparing results was evaluated by means of the Student’s *t* test.

## AUTHORS’ CONTRIBUTIONS

G.D.R. and P.I.N. carried out the genetic manipulations, quantitative physiology experiments, and *in vitro* enzyme assays. G.D.R., V.D.L., and P.I.N. conceived the whole study, designed the experiments, contributed to the discussion of the research and interpretation of the data, and wrote the article.

## ACKNOWLEDGMENTS

The authors are indebted to B. Calles (CNB-CSIC, Spain), M. H. Nørholm (Technical University of Denmark, Denmark), and A. Sinskey (Massachusetts Institute of Technology, USA) for helpful discussions and for sharing research materials. This study was supported by The Novo Nordisk Foundation (Grant NNF10CC1016517) and the Danish Council for Independent Research (*SWEET*, DFF-Research Project 8021-00039B) to P.I.N. This study was also supported by the *HELIOS* Project of the Spanish Ministry of Economy and Competitiveness BIO2015-66960-C3-2-R (MINECO/FEDER), and the *ARISYS* (ERC-2012-ADG-322797), *EmPowerPutida* (EUH2020-BIOTEC-2014-2015-6335536), and *MADONNA* (H2020-FET-OPEN-RIA-2017-1-766975) contracts of the European Union to V.D.L.

